# Quantitative characterization of translational riboregulators using an in vitro transcription-translation system

**DOI:** 10.1101/290403

**Authors:** Anis Senoussi, Jonathan Lee Tin Wah, Yoshihiro Shimizu, Jérôme Robert, Alfonso Jaramillo, Sven Findeiss, Ilka M. Axmann, André Estevez-Torres

## Abstract

Riboregulators are short RNA sequences that, upon binding to a ligand, change their secondary structure and influence the expression rate of a downstream gene. They constitute an attractive alternative to transcription factors for building synthetic gene regulatory networks because they can be engineered de novo and they have a fast turnover and a low metabolic burden. However, riboregulators are generally designed in silico and tested in vivo, which only provides a yes/no evaluation of their performances, thus hindering the improvement of design algorithms. Here we show that a cell-free transcription-translation (TX-TL) system provides valuable quantitative information about the performances of in silico designed riboregulators. In particular, we use the ribosome as an exquisite molecular machine that detects functional riboregulators, precisely measures their concentration and linearly amplifies the signal by generating a fluorescent protein. We apply this method to characterize two types of translational riboregulators composed of a cis-repressed (cr) and a trans-activating (ta) strand. At the DNA level we demonstrate that high concentrations of taDNA poisoned the activator until total shut off. At the RNA level, we show that this approach provides a fast and simple way to measure dissociation constants of functional riboregulators, in contrast to standard mobility-shift assays. Our method opens the route for using cell-free TX-TL systems for the quantitative characterization of functional riboregulators in order to improve their design in silico.

## 1 Introduction

During the early wave of synthetic biology (*1, 2*), known transcription factors were wired to their corresponding promoter sequences to control the expression of other transcription factors or effector proteins. While this approach has been very successful in engineering gene regulatory networks (GRNs) (*3*) with few nodes, the number of different elements in synthetic GRNs has stagnated at 5-6 (*4*). Two arguments may explain this limit. First, protein-DNA interactions are very difficult to design, although very promising computation methods are arising (*5*); the engineer must thus choose well-known transcription factor-promoter pairs. Second, the expression of these transcription factors imposes a metabolic burden to the cells (*6*).

Implementing regulatory circuits at the RNA level may help solving these issues because RNA-RNA interactions can be predicted from the sequence (*7* –*9*) and protein expression is not needed for regulation, which lowers the metabolic burden. The principal component of an RNA-regulated GRN is the riboregulator: an RNA sequence in the 5’ untranslated region (UTR) of a gene of interest that has an effect on its expression rate. Since they were first used in synthetic biology more than a decade ago (*10*), several riboregulators have been designed and implemented in vivo, both in procaryotic (*11* –*15*) and eukaryotic cells (*16*). However, their design remains more difficult than expected and many implementations do not work in vivo (*17*). One reason to this is that structure-prediction tools do not yet precisely capture the complexity involved in the folding of RNA species several hundreds of nucleotides long. In silico design needs furthermore a structural model on how regulation should work, which needs to be transformed into predictable features in order to generate optimized sequences. Another reason is that it is hard to control and tune the copy number of plasmids or genes in vivo and thus testing new parts in vivo (*18, 19*) often provides a yes/no answer that is difficult to correlate with thermodynamical parameters used in silico.

Including a phase of in vitro testing in the workflow of engineering riboregulators could potentially solve these problems. Structural characterization of riboregulators (*20, 21*) helps assessing the correctness of the designed structures but does not provide functional information and often involves complex experimental procedures. To overcome these difficulties and accelerate the improvement of *in-silico* designs, cell-free transcription-translation (TX-TL) platforms are an attractive tool for testing genetic regulatory modules in synthetic biology (*22, 23*). First, TX-TL in vitro testing can be used to qualitatively evaluate the performances of new designs in a faster manner, as it has been recently proposed (*23*). Second, it can provide quantitative data such as thermodynamic and kinetic rates that are of great value to improve in silico methods.

Here we used a purified TX-TL platform to illustrate the second approach. Its main advantage is that it uses the ribosome as an exquisite molecular machine that detects and amplifies the signal of functional riboregulators with great specificity, without making an a priori hypothesis about the structure of functional regulators. Importantly, we characterize the riboregulator dynamics at the DNA and RNA level, which allows to independently study transcription and translation and clearly pinpoint design shortcomings. Finally, our method provides dissociation constants of translational riboregulators that may help improving in silico design routines.

## 2 Results and discussion

### 2.1 Translation rate vs. structure as the optimization goal for a riboregulator

Our study focuses on translational riboregulators, which are composed of two RNA strands (Figure 1A). One of them, called cis repressed RNA, noted R_*cr*_, about 800 nucleotides (nt) long, codes for a gene but bears a hairpin in its 5’-untranslated region (5’-UTR) that prevents the ribosome to start reading the downstream gene. The other one, a small trans-activating RNA, about 100 nt long, noted R_*ta*_, hybridizes to the 5’-UTR of R_*cr*_, opens up the hairpin and forms an active complex, R_*act*_, which translation rate is increased.

**Figure 1:**
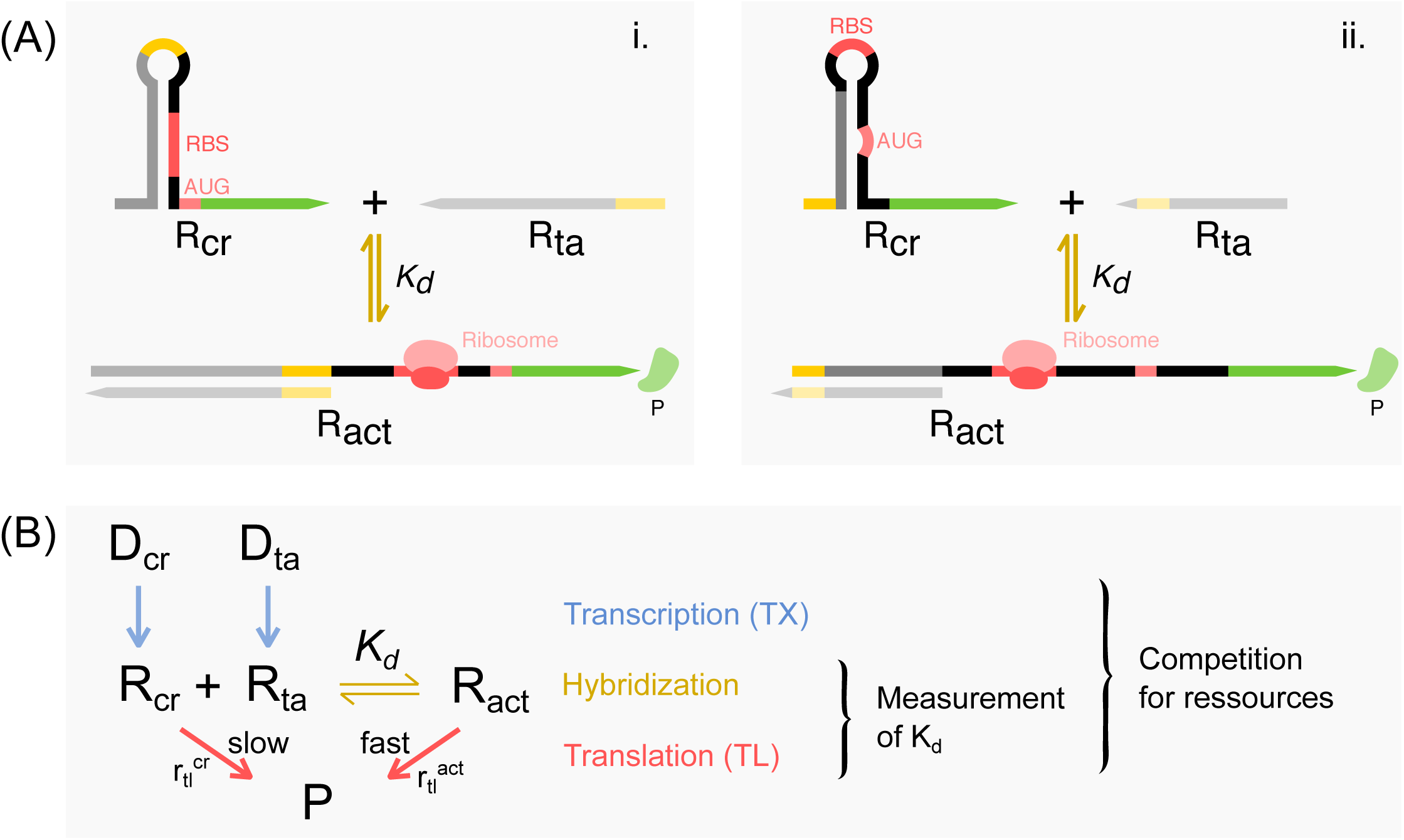
Principle of a transational riboregulator and of its characterization using a cell-free transcription-translation system (TX-TL). (A) Sketches of the two operation modes of translational riboregulators functioning as an activator. The 5’-UTR of R_*cr*_ RNA, forms a hairpin that hides either the ribosome binding site (RBS, i.) or the start codon (AUG, ii.) away from the ribosome. R_*ta*_ hybridizes with R_*cr*_, unwinding the hairpin and liberating the RBS and/or the AUG promoting translation. (B) Mechanism of transcription, riboregulation through RNA hybdridization and translation used in this work. DNA sequences D_*cr*_ and D_*ta*_ are transcribed into a cis-repressed, R_*cr*_, and a trans-activator, R_*ta*_, RNA strands. R_*cr*_ may be slowly translated into protein P or hybridize with R_*ta*_ to form R_*act*_ that is translated more rapidly into P. Measuring the dynamics of fluorescence production by a fluorescent protein P provides information about ressource competition when evaluating the system at the DNA level and quantitative values of dissociation constants *K*_*d*_ when RNA concentration is fixed.

Ultimately, the riboregulator engineer is interested in controlling the rate of translation for R_*cr*_ and R_*act*_, noted respectively *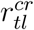* and *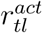*, and seek the objective *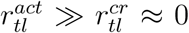* for an activator (Figure 1B). For convenience we assign a species name to an RNA sequence, but one must bear in mind that a given RNA sequence, for instance R_*cr*_, may fold in an ensemble of different structures *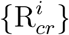*, with different translation rates *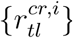*. Current in silico design methods (*9, 24*) compute the ensemble of secondary structures *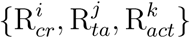* that minimizes free energy. However, the structure-to-function relationship that associates an RNA conformation with its translation rate is hard to establish. Thus, a set of heuristic rules attributes low values of translation rates *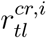* to structures where the RBS or the start codon are buried in a hairpin (Figure 1), and high values of *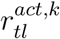*, to structures where these are accessible. However, these heuristic rules often fail. Moreover, minimizing the free energy of the RNA structures implies that the hypothesis of thermodynamic equilibrium holds, which is far from being true in vivo where co-transcriptional folding and RNA chaperones are the rule (*25, 26*).

To shed light into this problem we measured translation rates of recently in silico designed riboregulators (*18, 19*). Considering the difficulty of measuring translation rates in vivo, in particular because it is hard to control the equilibrium concentrations *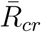* and 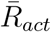, we used a cell-free TX-TL system called PURE system (Protein synthesis Using Recombinant Elements) (*27*). The PURE is composed of purified recombinant elements necessary for transcribing and translating a coding DNA or RNA sequence, totally in vitro. Briefly, the PURE system includes T7 RNA polymerase, an energy-coupling module for NTP regeneration, transfer RNAs, ribosomes, translation initiation, elongation and release factors in a suitable buffer (*28, 29*). By its intrinsic flexibility, a TX-TL system allows us to precisely tune the relative concentrations of the two components of a riboregulator at the DNA and at the RNA level. Moreover, the PURE system contains low level of ribonucleases, which is an essential property for having a reproducible and easily modelisable system (*30*).

### 2.2 The TX-TL system linearly amplifies the concentration of active RNA

We first characterized the translation and expression (transcription and translation) reactions of the PURE system in the absence of riboregulation. To do so, we prepared by PCR a linear DNA fragment coding for a green fluorescent protein (GFP) with no upstream regulatory region, called cr^−^DNA. It is composed of a T7 RNA polymerase promoter, a ribosome binding site and the GFP-coding sequence. To simplify transcription termination we did not add a terminator site at the end of the linear DNA fragment. In addition, we prepared by in vitro transcription the corresponding messenger RNA, cr^−^RNA, from cr^−^DNA. We successively used cr^−^RNA and cr^−^DNA as the coding nucleic acid input of the TX-TL system. We varied the concentration of the input and we measured the fluorescence emitted by the GFP produced over time (Figure 2). Starting from cr^−^RNA, the translation module of the TX-TL system actively produced GFP during 2 hours. The translation kinetics displayed three different phases: during about 5 min no signal was discernable from the background level, then followed a phase of quasi-linear increase during 100 min, that slowed down until a plateau was reached (Figure 2A). In the range 0 − 80 nM of cr^−^RNA, both the final intensity and the maximum rate of fluorescence growth, *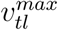* increased linearly with the initial quantity of coding RNA (Figure 2B). For higher concentrations there was a saturation: putting more RNA template did not increase significantly the final yield or the maximal production rate. When using cr^−^DNA as the initial input, the dynamics of the fluorescence intensity showed both common and contrasting features with the previous case (Figure 2C). Three phases were still observed: delay, growth and a plateau. However, the delay observed before an increase of fluorescence was now of 15 min. Finally, the quantity of DNA required to saturate the maximum rate of fluorescence growth, *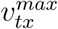*, was almost two orders of magnitude lower than the quantity of RNA that saturated translation (Figure 2D).

**Figure 2:**
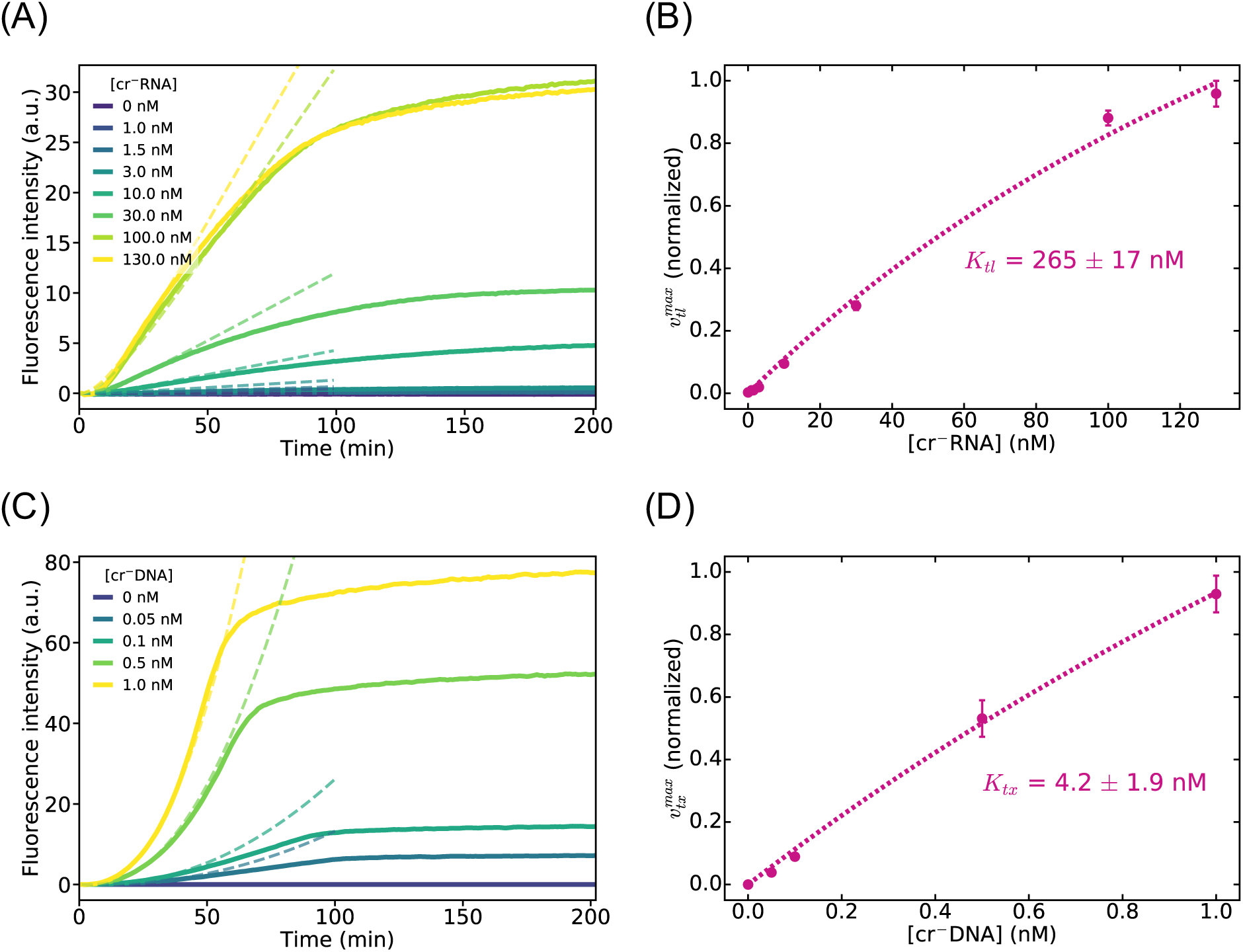
Characterization of the TX-TL system in the absence of riboregulation. Translation dynamics (A) and maximum fluorescence production rate (B) for increasing concentrations of an unregulated mRNA fragment coding for GFP. Expression (transcription and translation) dynamics (C) and maximum fluorescence production rate (D) for increasing concentrations of an unregulated linear DNA fragment coding for GFP. Solid lines (A,C) and disks (B,D) represent data, dotted lines are fits to the model. Error bars correspond to one sigma of a triplicate experiment.

We propose a simple quantitative kinetic model that fits our data. To take into account the saturation of the production rates we assigned Michaelis-Menten kinetics to the transcription and the translation reactions. As a plausible source of the initial delay in the translation reaction, we included a first-order step of maturation of the non-fluorescent GFP protein, noted P, into the functional fluorescent protein P*. This is in accordance to published maturation times (*31*). We neglected DNA and RNA degradation and we did not take into consideration the depletion of ressources because we analyzed our data between 0 and 50 min. For these reasons, our model did not reach a plateau in P* concentration (Figure 2A-C and Figure S2). These approximations are valid as long as the RNA molecules do not deteriorate and the enzymatic ressources, more specifically the ribosomes, are not depleted. We thus write the following mechanism

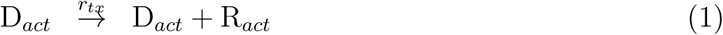

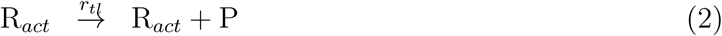

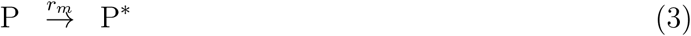

where D_*act*_ and R_*act*_ are, respectively, cr^−^DNA and cr^−^RNA and *r*_*tx*_, *r*_*tl*_ and *r*_*m*_ are, respectively, the transcription, translation and maturation rates. With the aforementioned hypotheses, this mechanism is associated with the rate equations

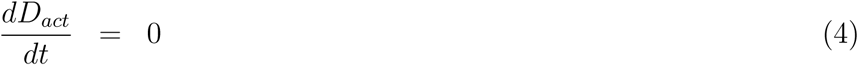

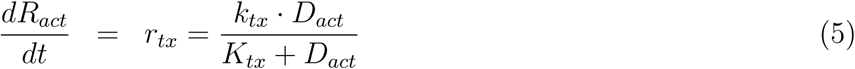

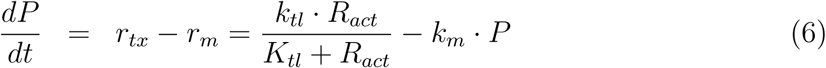

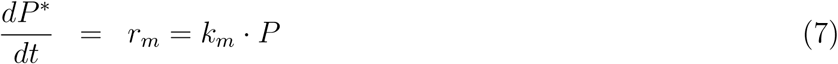

where *k*_*x*_ and *K*_*x*_ are, respectively, the rate and the Michaelis-Menten constants of reaction *x* and species concentrations are noted in italics. Equations (4-7) have exact solutions both for initial conditions corresponding to the translation (*D*_*act*_(0) = 0, *R*_*act*_(0) ≠ 0) and expression experiments (*D*_*act*_(0) ≠ 0, *R*_*act*_(0) = 0) (SI Section 4). For translation we obtain (SI Section 3.1)

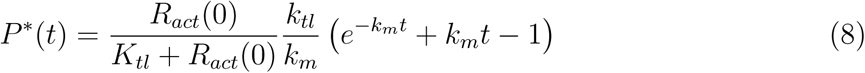

Note that when the ribosome is not saturated, *R*_*act*_(0) ≪ *K*_*tl*_, we can define a function *c*(*k*_*tl*_,*k*_*m*_,*t*) that does not depend on *R*_*act*_(0) and write

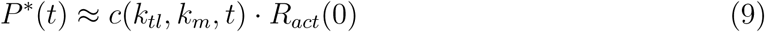

explicitly showing that translation acts as a linear amplifier of the initial concentration of active RNA. For expression, the exact solution is given in SI Section 3.3, here we provide an approximated solution when *R*_*act*_(*t*) ≪ *K*_*tl*_ (SI Section 3.2),

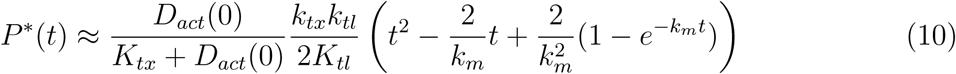

Considering that the fluorescence intensity is proportional to *P*\* we fitted (8) and (10) to the data in Figure 2. We obtained *K*_*tx*_ = 4.2 ± 1.9 nM, *K*_*tl*_ = 265 ± 17 nM and *k*_*m*_ = 0.10 ± 0.01 min^−1^, in fair agreement with previous measurements reporting *K*_*tx*_ = 4 − 9 nM for T7 RNA polymerase (*30, 32, 33*), *K*_*tl*_ = 66 nM (*30*) and *k*_*m*_ = 0.2 min^−1^ (*30, 34*). In summary, the saturation of transcription by DNA occurs at a concentration two-orders of magnitude lower than the saturation of translation by RNA and the TX-TL system acts as a linear amplifier of the concentration of active RNA, R_*act*_, with a readout of intensity fluorescence. As a result we can use GFP fluorescence as a measure of the concentration of R_*act*_.

### 2.3 Expression from DNA provides information on the saturation of transcriptional ressources

When riboregulators are used in vivo the DNA sequences D_*cr*_ and D_*ta*_, respectively coding for the cis-repressed and trans-activator RNA sequences R_*cr*_ and R_*ta*_, can either be inserted in the chromosome, in the same plasmid or in two different plasmids. This last case is common (*18, 35*) and a usual strategy to try to improve the riboregulator’s performance is to increase the effective concentration of R_*ta*_ by inserting D_*ta*_ in a high-copy plasmid. To test the effect of an increase in D_*ta*_ concentration in the performances of the riboregulator we performed a TX-TL expression experiment with two linear DNA fragments, D_*cr*_ and D_*ta*_. Within the TX-TL system the two DNA molecules are transcribed into the corresponding RNA strands, which associate into a coding RNA, R_*act*_. The production of P mainly comes from the translation of R_*act*_ but also may come from R_*cr*_, when cis-repression is not very effective. We thus write the following mechanism,

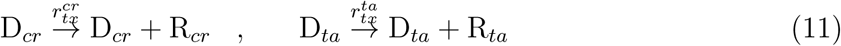

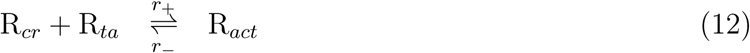

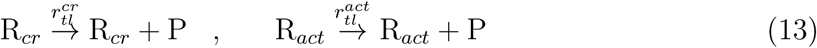

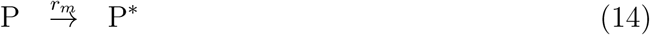

We titrated riboregulator G03 (SI Table S1) by keeping *D*_*cr*_ = 0.25 nM constant, varying *D*_*ta*_ in the range 0 − 100 nM and recording the GFP fluorescence over time (Figure 3). Increasing *D*_*ta*_ in the range 0 − 5 nM resulted in an increased fluorescence signal. However, for *D*_*ta*_ > 5 nM the fluorescence signal dramatically decreased until reaching 10% of the maximum production rate at *D*_*ta*_ = 100 nM. To explain this behaviour we hypothesized that D_*ta*_ and D_*cr*_ compete for transcriptional resources, i.e. a very high concentration of D_*ta*_ inhibits the transcription of D_*cr*_, thus reducing the concentration of R_*act*_. To test this hypothesis we titrated cr^−^DNA, which lacks the cis-regulatory region, with the D_*ta*_ of riboregulator G03. In agreement with our hypothesis, increasing D_*ta*_ steadily decreased the maximum fluorescence rate, *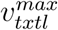* (Figure 3B), thus showing that the transcription of an orthogonal RNA strongly reduces the expression of the target mRNA.

**Figure 3:**
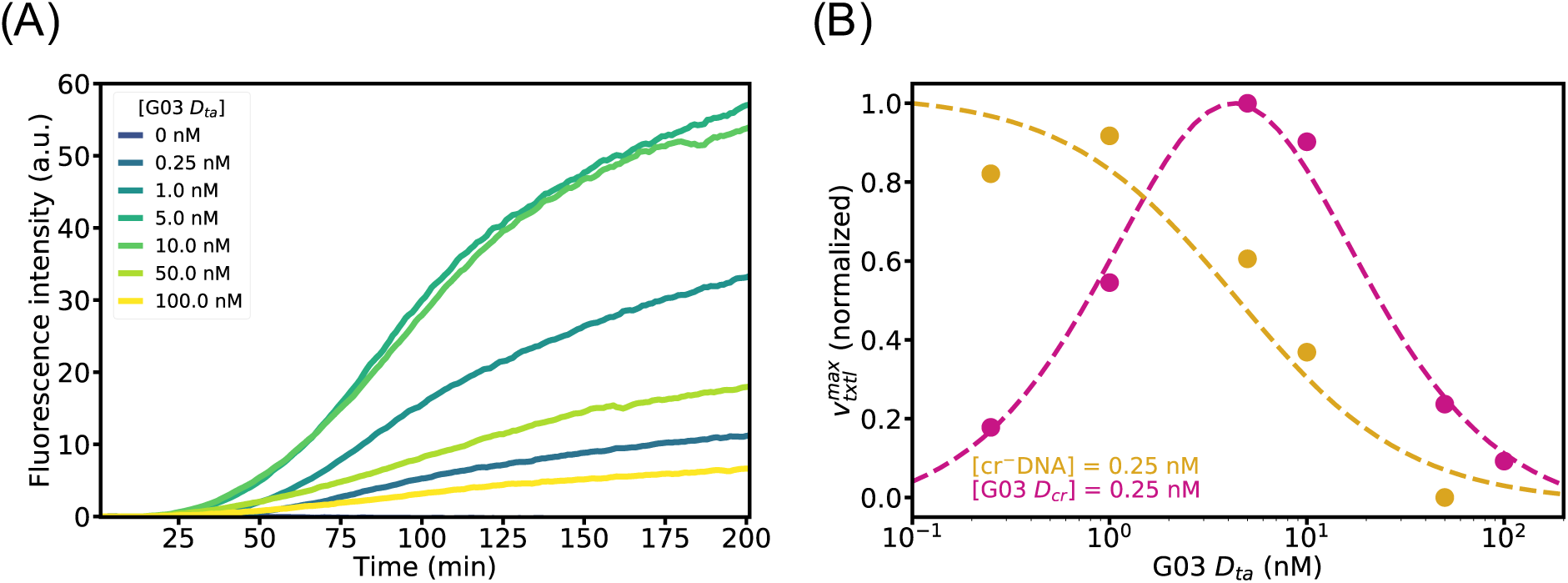
Titration of a riboregulator at the DNA level shows saturation of transcriptional ressources. (A) Fluorescence intensity vs. time for the in vitro expression of 0.25 nM of D_*cr*_ DNA, coding for GFP, with increasing concentrations of D_*ta*_ DNA, for riboregulator G03. (B) Normalized maximum fluorescence intensity production rate for a D_*cr*_ with, G03 (pink disks), or without, cr^−^ (yellow disks), cis regulatory region as a function of the concentration of D_*ta*_ from riboregulator G03. Dotted lines correspond to simulations of (6-7) together with (15-17).

To understand the role of saturation of transcriptional ressources, we modeled reactions (11-14) by the rate equations (4-7) but we replaced the production rate of R_*act*_ (5) by the following set of equations, that takes into account the competition for transcriptional ressources,

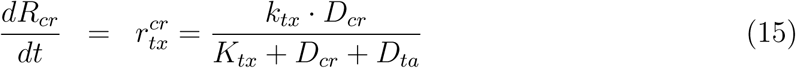

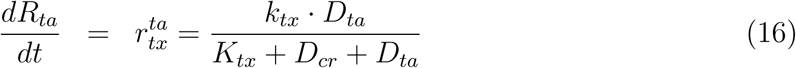

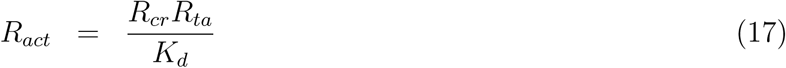

where we have assumed that the hybridization reaction (12), with dissociation equilibrium constant *K*_*d*_, is fast compared to the other reactions. From (15) it appears that *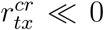* when *D*_*ta*_*/K*_*tx*_ ≫ 1, which we confirmed by solving the system of equations (6-7) together with (15-17), taking *K*_*d*_ = 100 nM (Table 1). We obtained the dashed lines in Figure 3 that are in very good agreement with experimental data. This in vitro result let us predict that inserting D_*ta*_ in a high-copy plasmid will decrease the performance of the riboregulator activator.

**Table 1:** Dissociation constants *K*_*d*_ at 37°C for the studied riboregulator devices measured using the cell-free translation method (txtl) and the mobility-shift method (ms). N.M. indicates that the electropherogram showed ill-defined peaks from which *K*_*d*_ could not be extracted.

| Device | $K_d^{txtl}$ (nM) | $K_d^{ms}$ (nM) |
| --- | --- | --- |
| RAJ11 | $15 \pm 14$ | N.M. |
| RAJ12 | $2220 \pm 950$ | N.M. |
| G01 | $46 \pm 56$ | $180 \pm 20$ |
| G03 | $134 \pm 80$ | $240 \pm 110$ |
| G80 | $31 \pm 19$ | $110 \pm 90$ |

### 2.4 Translation from RNA characterizes the reaction between the cis-repressed and the trans-activator RNA

The regulatory step of translational riboregulators takes place when the two RNA fragments, R_*cr*_ and R_*ta*_ hybridize and thereby change the accessibility of the ribosome to a site needed for initiating translation (RBS or AUG). The core of the riboregulation process can thus be described with reactions (12) and (13), where the first reaction involves the hybridization of R_*cr*_ with R_*ta*_ to form an active RNA complex, R_*act*_, that can be translated, and the second reaction the translation of R_*act*_ into protein P. It is not straightforward to characterize the thermodynamics of the first reaction. One possibility is to use an electrophoretic mobility shift assay in a polyacrylamide gel. Another way uses the property of a reverse transcriptase to terminate on stable RNA duplexes (*10*). In both cases these assays characterize the species R_*act*_ for being a duplex RNA but they are not sensitive to its translational activity. Here, instead, we probed the equilibrium concentration of R_*ta*_ that is active for translation. Our method is thus more meaningful to evaluate the design performances of a riboregulator.

We tested two types of riboregulators, two loop-mediated (*19*) and three toehold-mediated riboregulators (*18*). In the former, the RBS is buried inside the hairpin and the R_*ta*_ binds first to the loop on the hairpin. In the later, the start codon is protected by the hairpin and the R_*ta*_ binds to a toehold sequence on the 5’ side of the hairpin. We in vitro transcribed the R_*cr*_ and R_*ta*_ of these riboregulators (Figure S??) and studied their translation dynamics by titrating 5 nM R_*cr*_ with increasing concentrations of its corresponding R_*ta*_ in the range 0 − 1000 nM (Figure 4 and SI Figure S4). Because translation linearly amplifies *R*_*act*_ (Figure 2B and (9)), measuring the GFP intensity at a given time is directly proportional to the concentration of R_*act*_ that is translationally active. We thus plotted the normalized GFP fluorescence at 200 min as a function of the log of R_*ta*_ concentration. For a bimolecular equilibrium such as (12) one expects these plots to be described by

**Figure 4:**
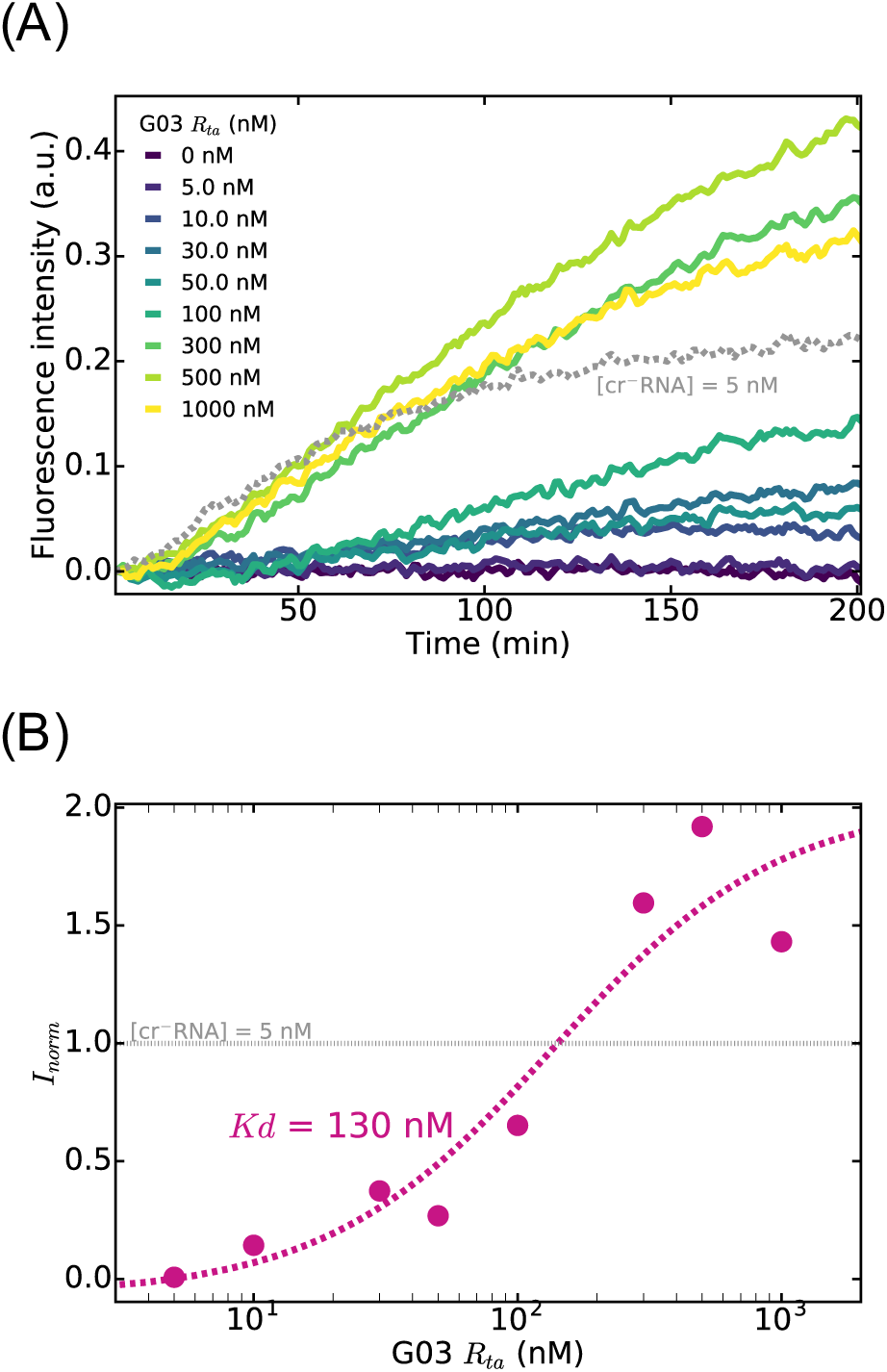
Titration of a riboregulator at the RNA level measures the dissociation constant of the riboregulator complex. GFP fluorescence produced over time (A) and normalized maximum fluorescence production rate (B) for different trans-activator concentrations, *R*_*ta*_. As a control, panel A shows the fluorescence intensity produced by the translation of 5 nM of an unregulated cr^−^RNA (orange dashes). In (B) disks correspond to experimental data and the dashed line is a fit of (18) to the data. Data for riboregulator G03.

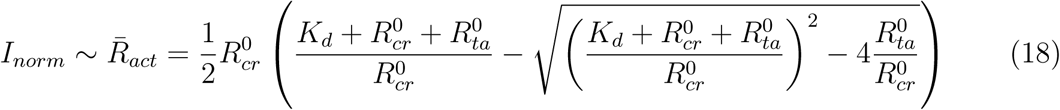

where *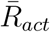* is the equilibrium concentration of R_*act*_ and superscript 0 indicates initial concentrations. Our experimental data followed well this trend (Figures 4 and S4). We thus fitted (18) to the data and found dissociation equilibrium constants in the range 10 − 2000 nM (Table 1), in agreement with *K*_*d*_ values of the order of 100 nM that have already been reported for loop-mediated activators (*10*). However, in one case, for G01, after a normal sigmoidal increase of *I*_*norm*_ vs. *R*_*ta*_, *I*_*norm*_ decreased for *R*_*ta*_ > 200 nM (Figure S4). To evaluate why in this particular case high *R*_*ta*_ inhibited translation, we performed a control experiment where a well-behaved regulator, G80, activated with 100 nM of its corresponding R_*ta*_, was titrated with increasing concentrations of R_*ta*_-G01 (Figure 5). We observed again that very high concentrations of R_*ta*_-G01 significantly reduced the final GFP concentration. We thus concluded that R_*ta*_-G01 poisoned the translation machinery, probably by nonspecific binding to other RNA components, including tRNAs, ribosomes, mRNAs, with about 1 *µ*M affinity.

**Figure 5:**
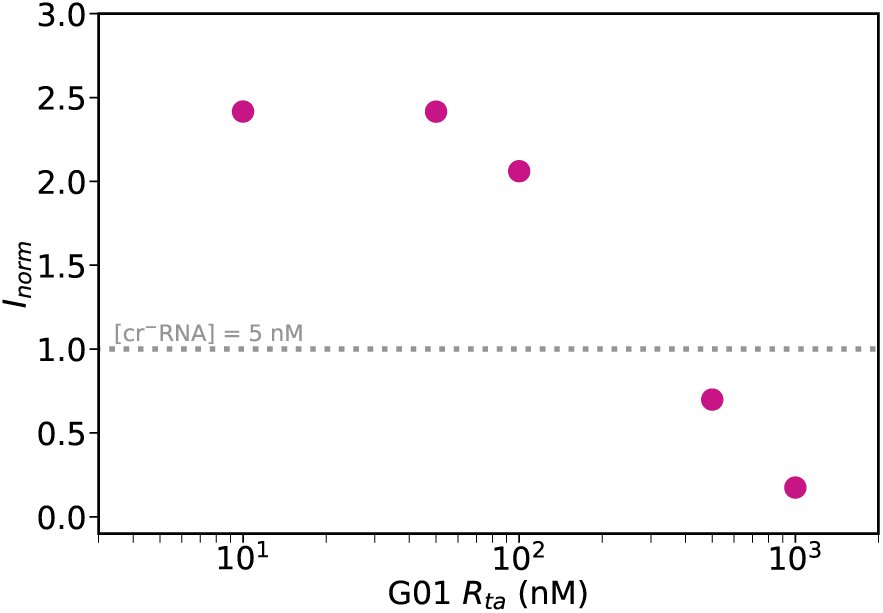
Titration of activated riboregulator G80 (*R*_*cr*_ = 5 nM *R*_*ta*_ = 50 nM) with increasing concentrations of R_*ta*_ from riboregulator G01.

To assess the performance of our method for measuring *K*_*d*_, we independently measured it with a standard mobility-shift assay performed with capillary gel electrophoresis. We used the same purified R_*cr*_ and R_*ta*_ that we mixed together at 37°C in a buffer with identical salt composition than the TX-TL system during 10 min before performing the electrophoresis assay. R_*cr*_ concentration was 8.3 nM and the R_*ta*_ concentration was ranging from 0 to 200 nM. Figure 6 shows the electropherograms for riboregulator G03, where a peak in intensity at a given time point corresponds to an RNA structure. In our experiments we detected three main peaks corresponding to R_*ta*_ at 22 s (Figure SS??) and R_*cr*_ and R_*act*_ complex between 37 and 40 s (Figure 6A). Interestingly, species R_*cr*_ and R_*act*_ yielded well-resolved peaks for toehold-mediated but not for loop-mediated riboregulators (Figure SS??). As a result this method only provided *K*_*d*_ for some but not all of the tested riboregulators, in contrast with the TX-TL method. The values obtained were of the same order of magnitude of those obtained by TX-TL. However, mobility-shift assay yielded *K*_*d*_ in a narrower range of 100 − 250 nM, while TX-TL was able to better discriminate *K*_*d*_ for the same species and provided values in the range 8 − 160 nM (Table 1).

**Figure 6:**
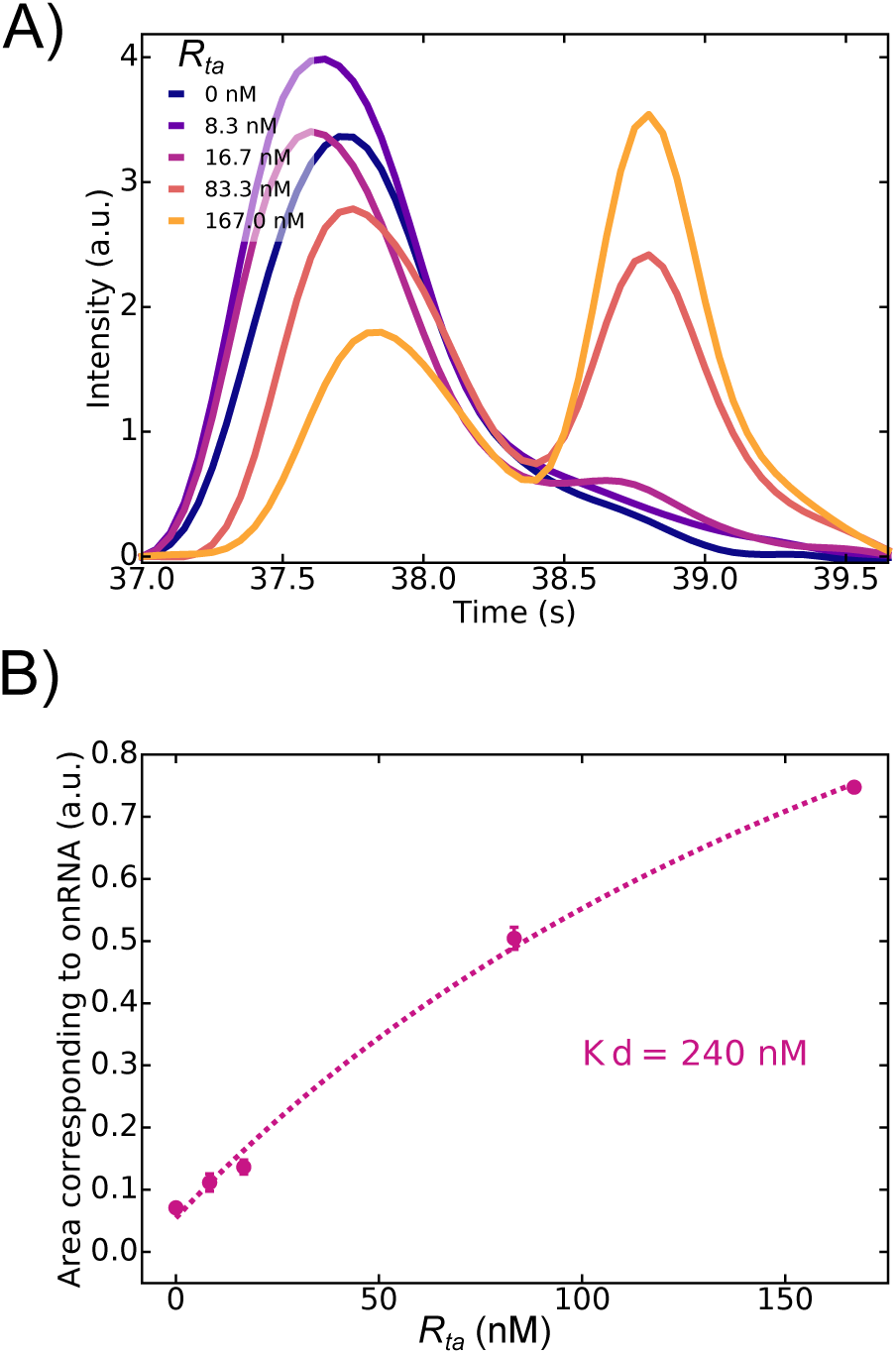
Titration of translational riboregulator G03 by mobility-shift capillary electrophoresis. (A) Corrected electropherograms vs. elution time and (B) peak area for different concentrations of R_*ta*_. Error bars correspond to one sigma of a triplicate experiment. Dashed line is a fit of (18) to the data.

## 3 Conclusion

We have demonstrated that in vitro transcription-translation (TX-TL) systems are an attractive platform to quantitatively characterize translational riboregulators. To do so we have taken advantage of the ribosome as a molecular machine that measures the concentration of RNA complexes that are translationally active. We have shown that increasing the DNA concentration of the trans-activating species inhibits expression by the saturation of the RNA polymerase, and we have predicted that inserting trans-activating elements in high-copy plasmids in vivo should limit the efficiency of translational activators. By titrating the cis-repressed gene with the trans-activating species at the RNA level we could determine dissociation constants, *K*_*d*_, for the RNA hybdridization reaction in a very simple manner. In particular, we could obtain *K*_*d*_’s for riboregulators that could not be resolved by mobility-shift assays. Our method thus provides a simple and rapid way for the quantitative characterization of riboregulators.

Combined with other biomolecular techniques such as molecular beacons (*33*) and automated-based designs (*36*), cell-free transcription-translation systems are becoming essential for a wide brand of applications. They allow to verify theoretical predictions on both RNA structures and behaviour of large scale regulatory networks. Their versality is a real asset for conceiving new synthetic biological features (*37*) and creating innovative biomolecular tools (*38*). The use of an in vitro step in the design and elaboration of complex synthetic regulatory networks will maximise the chance of expected in-vivo performances.

## 4 Methods

### DNA and RNA preparations

DNA templates were prepared by PCR amplification of plasmids encoding for the RNA translational regulators, followed by affinity column purification using Monarch PCR Purification Kit (New England BioLabs) or PureLink PCR Purification Kit (Thermo Fisher Scientific). Primers used for PCR amplification contained a T7 promotor or a T7 terminator (Biomers). RNA templates were prepared by in vitro transcription followed by purification using MEGAclear Transcription Clean-Up Kit (Ambion). The DNA and RNA integrity was determined by a 2% agarose gel and the concentrations were determined by absorbance at 260 nm using a NanoDrop 2000 UV-Vis spectrophotometer. The sequences of the riboregulator domains (Table S1), of the PCR primers (Table S2) and of the plasmids are compiled in the SI.

### Preparation of the PURE TX-TL system

The PURE TX-TL system was prepared according to (*39*) to reach the following composition: 1 units/*µ*L of RNase inhibitor Murine (New England Biolabs), 50 mM Hepes-KOH pH 7.6, 13 mM magnesium acetate, 100 mM potassium glutamate, 2 mM spermidine, 1 mM dithiothreitol (DTT), 2 mM of each ATP and GTP, 1 mM of each CTP and UTP, 20 mM creatine phosphate, 0.3 mM 20 amino acids, 56 A260/ml tRNA mix (Roche), 10 *µ*g/mL 10-formyl-5, 6, 7, 8-tetrahydrofolic acid, 0.1 mM each of amino acids, and factor mix. The factor mix contained 1.2 *µ*M ribosome, 10 *µ*g/ml IF1, 40 *µ*g/ml IF2, 10 *µ*g/ml IF3, 50 *µ*g/ml EF-G, 100 *µ*g/ml EF-Tu, 50 *µ*g/ml EF-Ts, 10 *µ*g/ml RF1, 10 *µ*g/ml RF2, 10 *µ*g/ml RF3, 10 *µ*g/ml RRF, 600-6000 U/ml of each ARS and MTF 4.0 *µ*g/ml creatine kinase (Roche), 3.0 *µ*g/ml myokinase (Sigma), 1.1 *µ*g/ml nucleoside-diphosphate kinase, 1.0 U/ml pyrophosphatase (Sigma), and 10 *µ*g/ml of T7 RNA polymerase.

### Fluorescence measurements in real-time PCR machine

Rotor-GeneQ real-time PCR (Qiagen) was used to record fluorescence from GFP expression (excitation 470±10 nm, emission 510± 5 nm) in a 15 *µ*L volume. The temperature was set to 37°C and fluorescence recorded every minute for at least 3 h.

### Data processing

Data were processed using in-house Python routines. For each condition of template — DNA or RNA— concentration, fluorescence intensity plots were shifted to the origin by removing the mean value of the three first minutes and by subtracting the fluorescence due to the PURE TX-TL system without any template. Corrected data were filtered using a Savitzky–Golay filter (window length: 21, polynomial order: 3) to remove residual noise before being derived to compute *v*^*max*^.

### Electrophoretic mobility shift assays

Electrophoretic mobility shift assays were performed with a 2100 Bioanalyzer System (Agilent Technologies) and an RNA Nano chip Kit. Samples were prepared by mixing RNA strands in 50 mM Hepes-KOH pH 7.6, 13 mM magnesium acetate, 100 mM potassium glutamate, 2 mM spermidine, 1 mM DTT and nuclase free water. They were incubated at 37°C for 10 min before being loaded into the electrophoresis chip. Electropherograms were manually aligned along the time axis. Affine curves corresponding to the backgrounds of zones of interest were subtracted. Areas under peaks were determined by numerical integration and were normalized using an RNA marker provided in Agilent’s kit.

## Acknowledgement

The authors thank H. Isambert for helpful discussions, A. Green for providing the expression plasmids coding for the toehold-mediated riboegulators and J.-C. Galas for comments on the manuscript. This research was supported by the European commission FET-Open program under award Ribonets (323987).

## Supporting Information Available

This material is available free of charge via the Internet at http://pubs.acs.org/.

