## Supplementary Materials for "Quantitative characterization of translational riboregulators using an in vitro transcription-translation system"

### Supplementary information for: Quantitative characterization of translational riboregulators using an in vitro transcription-translation system

Anis Senoussi,[†,⊥] Jonathan Lee Tin Wah,[†,⊥] Yoshihiro Shimizu,[‡] Jérôme Robert,[†] Alfonso Jaramillo,[¶,#] Sven Findeiss,[§,@] Ilka M. Axmann,[‖] and André Estevez-Torres[*,†,⊥]

†*Sorbonne Universités, UPMC Univ Paris 06, Laboratoire Jean Perrin, F-75005, Paris, France*

‡*Laboratory for Cell-Free Protein Synthesis, RIKEN Quantitative Biology Center, 6-2-3, Furuedai, Suita, Osaka 565-0874, Japan*

¶*School of Life Sciences and WISB, University of Warwick, Gibbet Hill Road, Coventry, CV4 7AL, United Kingdom*

§*Dept. Computer Science, and Interdisciplinary Center for Bioinformatics, University Leipzig, Härtelstrasse 16-18, D-04107 Leipzig, Germany*

‖*Institute for Synthetic Microbiology, Cluster of Excellence on Plant Sciences (CEPLAS), Heinrich Heine University Düsseldorf, Universitätsstrasse 1, 40225 Düsseldorf, Germany*

⊥*UMR 8237, CNRS, F-75005, Paris, France*

#*Institute of Systems and Synthetic Biology, Batiment Geneavenir 6, 5, rue Henri Desbrueres, 91030 Evry Cedex, France*

@*University of Vienna, Faculty of Computer Science, Research Group Bioinformatics and Computational Biology and Faculty of Chemistry, Department of Theoretical Chemistry, Währingerstrasse 29, A-1090 Vienna, Austria*

**4 Model for competition of transcriptional ressources into the PURE cell-free system** **9**

**5 Kd determination and equation** **10**

**6 RNA titration for other translational regulators using TX-TL** **11**

**7 Electrophoretic mobility shift assay data** **12**

**References** **12**

### 1 Sequences

#### 1.1 RNA regulators sequences

Table S1: RNA sequences of the regulators. $R_{cr}$ sequences are listed from the end of the T7 promotor to the end of the regulating part. The ribosome binding site (RBS) is indicated in blue, the start codon in pink. $R_{ta}$ sequences are listed from the end of the T7 promotor and before the beginning of the T7 terminator.

| **Regulator** | **Sequences 5' → 3'** |
|---|---|
| $cr^-$ RNA | AAAGAGGAGAAA UUAUGA AUG |
| RAJ11 $R_{cr}$ | CUCGCAUAAUUUCACUUCUUCAAUCCUCCCGUU AAAGAGGAGAAA UUAUGA AUG |
| RAJ11 $R_{ta}$ | GGGAGGGUUGAUUGUGUGAGUCUGUCACAGUUCAGCGGAAACGUUGAUGCUGUGACAGAUUUAU-GCGAGGC |
| RAJ12 $R_{cr}$ | ACCCAGUAUCAUUCUCUUCUUCCUGCCCACGCGG AAAGAGGAGAAA GGUGUA AUG |
| RAJ12 $R_{ta}$ | GGGCAGGAAGAAGGGUUCCUUUGAGCGAAUCUAGCGGCACCUCGCUAGGAUUUGCUCGAAGGGA-UUCUGGG |
| G01 $R_{cr}$ | GGGUGAAUGAAUUGUAGGCUUGUUAUAGUUAUGAAC AGAGGAGA CAUAAC AUG AACAAGCCU |
| G01 $R_{ta}$ | GGGACCGUGGACCGCAUGAGGUCCACGGUAAACAUAACUAUAACAAGCCUACAAUUCAUUCAAAC |
| G03 $R_{cr}$ | GGGUGAUGGAAUAAGGCUGUGUAUAUGAUGUUAGAC AGAGGAGA UAACAU AUG AUACACAGC |
| G03 $R_{ta}$ | GGGUCAGUUCCUGAGGUACCAGGAACUGAAACUAACAUCAUAUACACAGCCUUAUUCCAUCACAC |
| G80L18 $R_{cr}$ | GGGUGAAUUUGAUUGACUAGAAUGAUGAUACGAAGACAAGAAC AGAGGAGA UCGUAU AUG CA-UUCUAGU |
| G80 $R_{ta}$ | GGGCCACGCGUUGUCCUAUCAACGCGUGGAAAUCGUAUCAUCAUUCUAGUCAAUCAAAUUCAAAG |

#### 1.2 Primers for PCR

Table S2: Primers for production of DNA fragments coding for translational regulators from plasmids. The plasmids coding for the Green regulators already contained T7 promotor and T7 terminator in their sequences. We used the two same primers for amplifying the Green regulators by PCR.

| **Primers** | **Sequences 5' → 3'** |
|---|---|
| RAJ11 crDNA FP | TAATACGACTCACTATAGGCCTCGCATAATTTCACTTCTTCAATCC |
| RAJ11 crDNA RP | CAAAAAACCCCTCAAGACCCGTTTAGAGGCCCCAAGGGGTTATGC-<br>TATTATTTGTATAGTTCATCCATGCCATGTGTAATC |
| RAJ11 taDNA FP | CAAAAAACCCCTCAAGACCCGTTTAGAGGCCCCAAGGGGTTATGC-<br>TAGCCTCGCATAAATCTGTCACAG |
| RAJ11 taDNA RP | TAATACGACTCACTATAGGGGGAGGGTTGATTGTGTGAGTC |
| RAJ12 crDNA FP | TAATACGACTCACTATAGGACCCAGTATCATTCTCTTCTTCCTGCC |
| RAJ12 crDNA RP | CAAAAAACCCCTCAAGACCCGTTTAGAGGCCCCAAGGGGTTATGC-<br>TATCATCATTTGTACAGTTCATCCATACCATGC |
| RAJ12 taDNA FP | ATCGTATTGGGGAACCCCGGAGATTTGCCCAGAACTCCCCAAAAA-<br>AGCCTCGCATAAATCTGTCACAGCATC |
| RAJ12 taDNA RP | TAATACGACTCACTATAGGGGGCAGGAAGAAGGGTTCCTTT |
| Green crDNA and taDNA FP | CGCGCTAATACGACTCACTATAGG |
| Green crDNA and taDNA RP | CAAAAAACCCCTCAAGACCCGTT |

#### 1.3 Plasmids

Plasmids coding for G01 $R_{cr}$, G01 $R_{ta}$, G03 $R_{cr}$, G03 $R_{ta}$ and G80 $R_{ta}$ are, as they appeared in (*1*), pAG_TS1_KS001, pAG_TS1_AT001, pAG_TS1_KS003, pAG_TS1_AT003 and pAG_TS1_AT080, respectively. Plasmid coding for G80L18 $R_{cr}$ is a variant of the pAG_TS1_KS080 plasmid.

Plasmids coding for RAJ11 and RAJ12 systems can be found on Addgene plasmid repository (Addgene plasmids #39244 and #3924, respectively).

### 2 Purity of the riboregulators

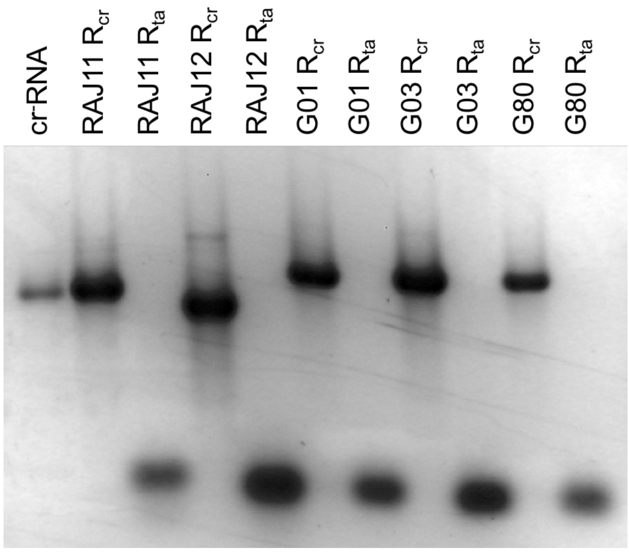

Figure S1: Denaturing 1.5% agarose gel of the in vitro transcribed riboregulators. Switch designates $R_{cr}$ and trigger $R_{ta}$.

### 3 Detailed solution of translation and expression kinetics in the absence of regulation

#### 3.1 Translation

From Eqs. (1-7) in the MT, translation corresponds to initial conditions ($D_{act}(0) = 0$, $R_{act}(0) = R^0_{act}$ and $P(0) = P^*(0) = 0$). We should thus consider the system of equations

$$\frac{dP}{dt} = r_{tx} - r_m = \kappa_{tl} - k_m \cdot P \quad (1)$$

$$\frac{dP^*}{dt} = r_m = k_m \cdot P \quad (2)$$

where we have noted $\kappa_{tl} = \frac{k_{tl} \cdot R^0_{act}}{K_{tl} + R^0_{act}}$. Calling the total protein concentration $P_{tot} = P + P^*$ and adding (1) and (2) we have

$$\frac{dP_{tot}}{dt} = \kappa_{tl} \quad (3)$$

which solution is $P_{tot}(t) = \kappa_{tl} t$. Combining (2) and (3) we have

$$\frac{dP^*}{dt} = k_m(\kappa_{tl} t - P^*) \tag{4}$$

that is a linear ODE that can be written in the form

$$g(t)y' = f_1(t)y + f_0(t), \tag{5}$$

with $g = 1$, $f_1 = -k_m$ and $f_0 = k_m \kappa_{tl} t$. Following reference (*2*), the solution of (5) is

$$y = Ce^F + e^F \int e^{-F} \frac{f_0(t)}{g(t)} dt, \text{where} \qquad F(t) = \int \frac{f_1(t)}{g(t)} dt \tag{6}$$

We find $F(t) = -k_m t$ and the solution of (4) is thus

$$P^* = \frac{\kappa_{tl}}{k_m} \left( e^{-k_m t} + k_m t - 1 \right) \tag{7}$$

which corresponds to (8) in the MT.

#### 3.2 Expression with non-saturated translation

From Eqs. (1-7) in the MT, expression corresponds to initial conditions $D_{act}(0) = D^0_{act}$, $R_{act}(0) = P(0) = P^*(0) = 0$. If we suppose that $R_{act}(t) \ll K_{tl}$ and we note $\kappa_{tx} = \frac{k_{tx} D^0_{act}}{K_{tx} + D^0_{act}}$ we should thus consider the system of equations

$$\frac{dR_{act}}{dt} = r_{tx} = \kappa_{tx} \tag{8}$$

$$\frac{dP}{dt} = r_{tx} - r_m \approx \frac{k_{tl}}{K_{tl}} - k_m \cdot P \tag{9}$$

$$\frac{dP^*}{dt} = r_m = k_m \cdot P \tag{10}$$

we find $R_{act} = \kappa_{tx} t$ and

$$P_{tot} \approx \frac{k_{tl}}{K_{tl}} \frac{\kappa_{tx}}{2} t^2 \tag{11}$$

and the variation of $P^*$ becomes

$$\frac{dP^*}{dt} \approx k_m \left( \frac{k_{tl}}{K_{tl}} \frac{\kappa_{tx}}{2} t^2 - P^* \right) \tag{12}$$

which is again a linear ODE. Comparing (12) with (5) we have $g = 1$, $f_0 = k_m \frac{k_{tl}}{K_{tl}} \frac{\kappa_{tx}}{2} t^2$ and $f_1 = -k_m$. We find again $F(t) = -k_m t$ and from (6) the solution of (12) is

$$P^*(t) \approx \frac{k_{tl}}{K_{tl}} \frac{\kappa_{tx}}{2} \left( t^2 - \frac{2}{k_m} t + \frac{2}{k_m^2} (1 - e^{-k_m t}) \right) \tag{13}$$

#### 3.3 Expression with Michaelis-Menten translation

We now consider the full system of equations (4-7) in the MT. The exact equation describing the variation of $P_{tot}$ is

$$\frac{dP_{tot}}{dt} = \frac{k_{tl} \kappa_{tx} t}{K_{tx} + \kappa_{tx} t} \tag{14}$$

which solution is

$$P_{tot} = k_{tl} \left( t - \frac{K_{tl}}{\kappa_{tx}} \ln \left( 1 + \frac{\kappa_{tx}}{K_{tl}} t \right) \right) \tag{15}$$

and we may check that a second order Taylor expansion of (15) is identical to (11), as expected. Combining (15) and (10) with the conservation of protein we have

$$\begin{aligned} \frac{dP^*}{dt} &= k_m (P_{tot} - P^*) && (16) \\ &= k_m \left[ k_{tl} \left( t - \frac{K_{tl}}{\kappa_{tx}} \ln \left( 1 + \frac{\kappa_{tx}}{K_{tl}} t \right) \right) - P^* \right] && (17) \end{aligned}$$

which is, once more, a linear ODE like (5) with $g = 1$, $f_1 = -k_m$ and $f_0 = k_m P_{tot}$. From (6) we obtain

$$P^* = Ce^{-k_m t} + e^{-k_m t} \int e^{k_m t} k_m \left[ k_{tl} \left( t - \frac{K_{tl}}{\kappa_{tx}} \ln \left( 1 + \frac{\kappa_{tx}}{K_{tl}} t \right) \right) \right] dt \tag{18}$$

Using the integrals

$$\int t e^{\alpha t} dt = \frac{\alpha t - 1}{\alpha^2} e^{\alpha t} \tag{19}$$

$$\int e^{\alpha t} \ln(t) dt = \frac{1}{\alpha} \left( e^{\alpha t} \ln |t| - \int_{-t}^{\infty} \frac{e^{-\alpha t}}{\alpha t} dt \right) \tag{20}$$

we find

$$P^* = Ce^{-k_m t} + \frac{k_{tl}}{k_m}(k_m t - 1) - \frac{k_{tl} K_{tl}}{k_m \kappa_{tx}} \ln \left| 1 + \frac{\kappa_{tx}}{K_{tl}} t \right| + e^{-k_m t} e^{-k_m K_{tl}/\kappa_{tx}} \int_{-1-\frac{\kappa_{tx}}{K_{tl}} t}^{\infty} \frac{e^{-u}}{u} du \tag{21}$$

with $C$ an integration constant.

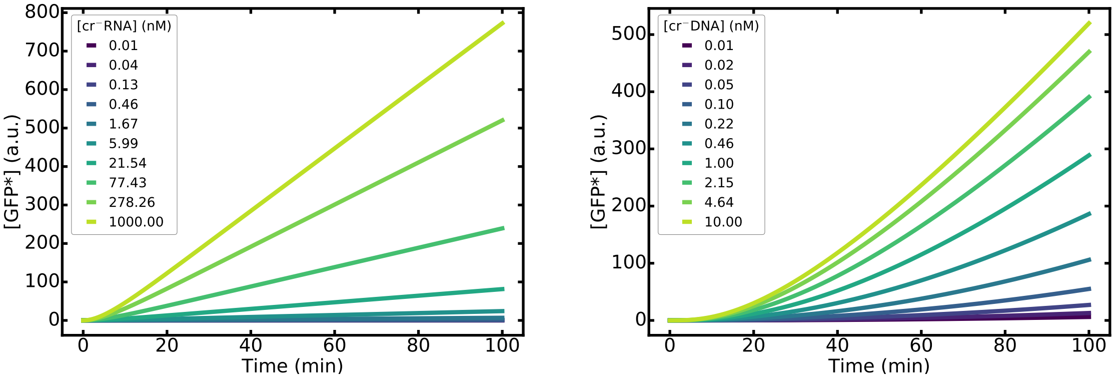

Figure S2: Simulation of the expression and the translation of the PURE system. (a) Translation from a coding RNA fragment. (b) Transcription and translation from a coding DNA fragment.

### 4 Model for competition of transcriptional ressources into the PURE cell-free system

We used the follow kinetic parameters in our model:

Table S3: Kinetic parameters for the study of RNA transregulation using the PURE cell-free system

| **Kinetic parameters** | |
|---|---|
| $k_{tx}$ (nM·min$^{-1}$) | 1 |
| $K_{TX}$ (nM) | 4 |
| $k_{tl}$ (nM·min$^{-1}$) | 1 |
| $K_{TL}$ (nM) | 265 |
| $K_d$ (nM) | 100 |
| $k_m$ (min$^{-1}$) | 0.1 |

The values for $K_{TX}$, $K_{TL}$ and $k_m$ were measured in this work (Figure 2). $k_{tx}$ and $k_{tl}$ were set to unity as they are irrelevant for the dynamics and only set the absolute concentration of P*, which we did not measure in our study (see Eqs. 8 and 10 in MT). $K_d$ was chosen to reach a typical value obtained in our work (Table 1).

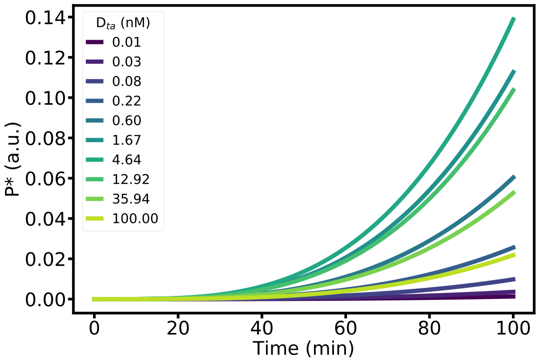

Figure S3: Simulation of the evolution of the P* concentration produced by a transregulated gene for different concentrations of $D_{ta}$.

### 5 Kd determination and equation

For the bimolecular reaction between $R_{cr}$ and $R_{ta}$ to give complex $R_{act}$ with initial concentrations of, respectively, $R^0_{cr}$, $R^0_{ta}$ and $R^0_{act} = 0$ we have

| $R_{cr}$ | + | $R_{ta}$ | = | $R_{act}$ |
|---|---|---|---|---|
| $R^0_{cr}$ | | $R^0_{ta}$ | | 0 |
| $R^0_{cr}(1 - X)$ | | $R^0_{ta}(1 - \frac{R^0_{cr}}{R^0_{ta}} X)$ | | $R^0_{cr} X$ |

where $X$ denotes the extent of the reaction.

The dissociation constant $K_d$ is defined by

$$K_d = \frac{\bar{R}_{cr} \bar{R}_{ta}}{\bar{R}_{act}} = \frac{R^0_{ta}(1 - \bar{X})(1 - \frac{R^0_{cr}}{R^0_{ta}} \bar{X})}{\bar{X}} \tag{22}$$

where bars denote equilibrium. Then, we obtain:

$$\bar{X}^2 - \frac{K_d + R^0_{cr} + R^0_{ta}}{R^0_{cr}} \bar{X} + \frac{R^0_{ta}}{R^0_{cr}} = 0. \tag{23}$$

The only non-negative solution of this second order equation is

$$\bar{X} = \frac{1}{2} \left( \frac{K_d + R^0_{cr} + R^0_{ta}}{R^0_{cr}} - \sqrt{\left( \frac{K_d + R^0_{cr} + R^0_{ta}}{R^0_{cr}} \right)^2 - 4 \frac{R^0_{ta}}{R^0_{cr}}} \right). \tag{24}$$

And thus we have

$$\bar{R}_{act} = R^0_{cr} \bar{X}. \tag{25}$$

### 6 RNA titration for other translational regulators using TX-TL

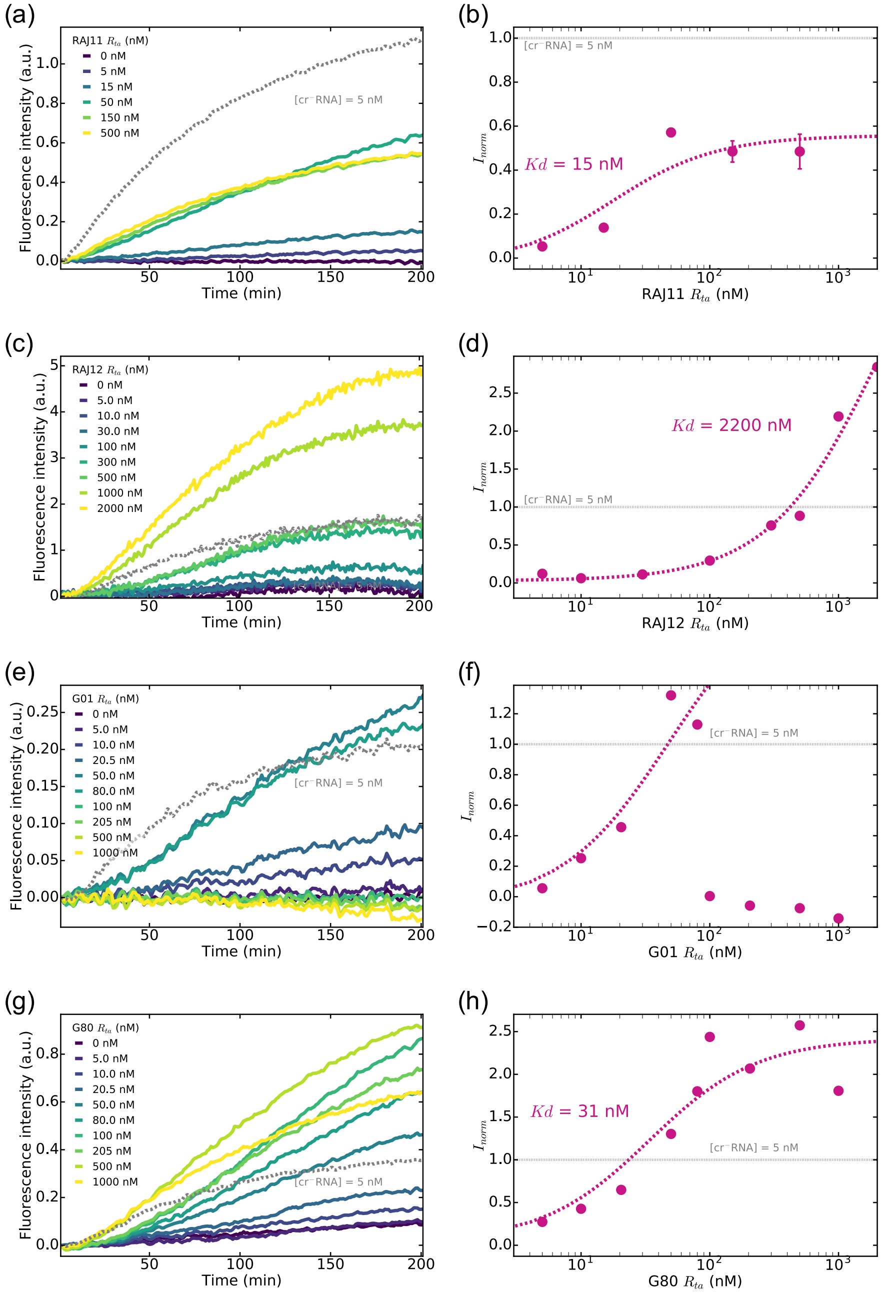

Figure S4: Titration of a translational riboregulator using the PURE cell-free system. Fluorescence from GFP produced over time (a,c,e,g) and normalized maximum fluorescence production rate for different initial $R_{ta}$ concentrations (b,d,f,h) for different riboregulators: RAJ11 (a,b), RAJ12 (c,d), G01 (e,f) and G80 (g,h) We used two controls to normalize titration plots: the intensity due to the PURE system mix without nucleic acids template (black dashes) and the fluorescence intensity produced by the translation of 5 nM of an unregulated gene RNA $cr^{-}$RAJ11 (grey dashes).

### 7 Electrophoretic mobility shift assay data

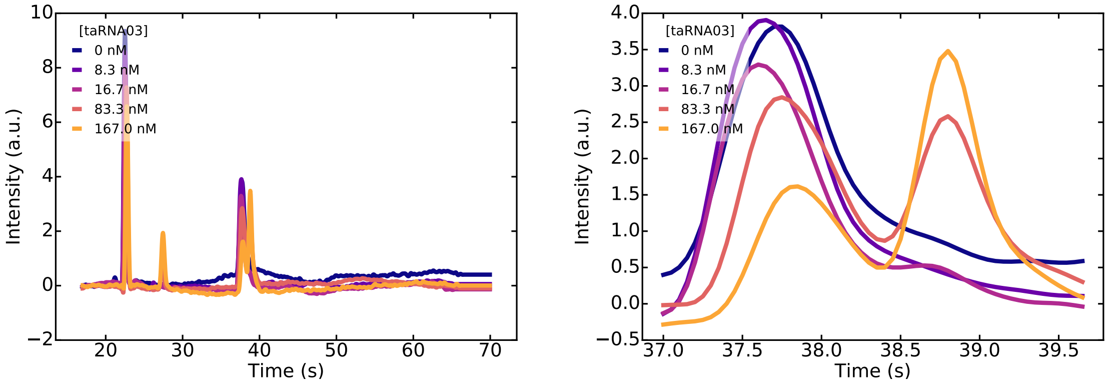

Figure S5: Effect of data processing on titration electropherograms. Raw (A) and corrected (C) data and zoom on raw (B) and corrected (D) data for riboregulator G03. The peak at 22 s corresponds to $R_{ta}$ and the peak at 28 s corresponds to an internal standard.

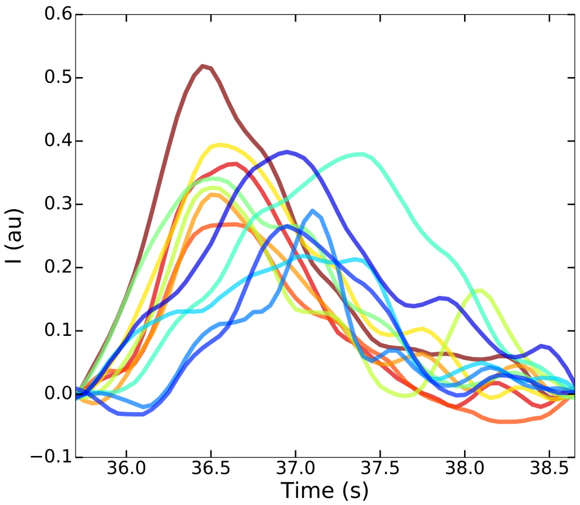

Figure S6: Corrected electrophoregrams vs. eluation time for the titration of translational riboregulator RAJ12 by mobility-shift capillary electrophoresis.
